# The frequency dependence of prestin-mediated fast electromotility for mammalian cochlear amplification

**DOI:** 10.1101/2024.05.22.595389

**Authors:** Satoe Takahashi, Yingjie Zhou, Mary Ann Cheatham, Kazuaki Homma

## Abstract

Prestin’s voltage-driven motor activity confers sound-elicited somatic electromotility in auditory outer hair cells (OHCs) and is essential for the exquisite sensitivity and frequency selectivity of mammalian hearing. Lack of prestin results in hearing threshold shifts across frequency, supporting the causal association of variants in the prestin-coding gene, *SLC26A5*, with human hearing loss, DFNB61. However, cochlear function can tolerate reductions in prestin-mediated OHC electromotility. We found that two deafness-associated prestin variants, p.A100T and p.P119S, do not deprive prestin of its fast motor function but significantly reduce membrane expression, leading to large reductions in OHC electromotility that were only ∼30% of wildtype (WT). Mice harboring these missense variants suffered congenital hearing loss that was worse at high frequencies; however, they retained WT-like auditory brainstem response thresholds at 8 kHz, which is processed at the apex of the mouse cochlea. This observation suggests the increasing importance of prestin-driven cochlear amplification at higher frequencies relevant to mammalian hearing. The observation also suggests the promising clinical possibility that small enhancements of OHC electromotility could significantly ameliorate DFNB61 hearing loss in human patients.

**SIGNIFICANCE:** Prestin is abundantly expressed in the auditory outer hair cells and is essential for normal cochlear operation. Hence, reduction of prestin expression is often taken as indicative of reduced cochlear function in diseased or aged ears. However, this assumption overlooks the fact that cochlear function can tolerate large reductions in prestin motor activity. DFNB61 mouse models generated and characterized in this study provide an opportunity to gauge the amount of prestin motor activity needed to sustain normal hearing sensitivity. This knowledge is crucial not only for understanding the pathogenic roles of deafness-associated variants that impair OHC electromotility but also for unraveling how prestin contributes to cochlear amplification.

## INTRODUCTION

*SLC26A5* encodes the homodimeric voltage-dependent membrane-based molecular motor, prestin (1-5), that is abundantly expressed in auditory outer hair cells (OHCs) in the cochlea. The voltage-driven motor activity of prestin confers sound-elicited somatic elongation and contraction, referred to as OHC electromotility (6). Mice lacking prestin (7-9) or expressing a virtually nonelectromotile artificial prestin mutant, p.V499G/p.Y501H prestin (10), suffer ∼50 dB threshold shifts across frequency. A recent study also showed that the p.R130S prestin variant, which was identified in human patients (11), slows prestin’s motor activity and induces frequency-dependent threshold elevations (worse at higher frequencies), indicating the mechanical contribution of prestin-mediated electromotility to cochlear amplification on a cycle-by-cycle basis (12). These studies established the essentiality of prestin for mammalian hearing and support the causal association of *SLC26A5* variants with autosomal recessive nonsyndromic sensorineural hearing loss, DFNB61 (OMIM: 613865). However, reductions in OHC electromotility by ∼50% barely or only slightly affect sensitivity (13-15). The mechanism underlying this counterintuitive tolerance remains enigmatic but points to the possibility of ameliorating DFNB61 hearing loss without restoring OHC electromotility to the wildtype (WT) level. Determining the relationship between the magnitude of OHC electromotility and hearing sensitivity is thus crucial not only for defining the cochlear amplification mechanism but also for assessing the possibility of clinical interventions (e.g., gene replacement therapies, pharmacological interventions, etc.) for DFNB61 hearing loss.

Naturally occurring prestin variants are invaluable for elucidating prestin’s physiological role. In this study, we characterized the functional consequences of two DFNB61-associated prestin variants identified in human patients with hearing loss, p.A100T (c.298G>A) and p.P119S (c. 355C>T) (16, 17), in mouse models. Although these missense variants do not deprive prestin of its fast motor function, they do significantly reduce membrane protein expression, resulting in hugely reduced OHC electromotility and hearing loss. However, despite the large reductions in OHC electromotility, sensitivity at frequencies coded at the apex of the mouse cochlea was near-normal, attesting to the resilience of cochlear function to reduced prestin motor activity.

## RESULTS

### The p.A100T and p.P119S prestin variants reduce the sound-induced mechanical response of OHCs

Recent cryo-EM studies provided significant structural and mechanistic insights for prestin (2-5). Ala^100^ and Pro^119^ are located in the transmembrane domain but do not directly contribute to the anion substrate binding (see Fig. 3 in Takahashi et al., 2024 (18)). These nonpolar residues are packed close to other nonpolar residues (Ala^100^ with Met^225^, Leu^440^, and Val^444^; Pro^119^ with Ile^134^, Cys^381^, and Ile^393^), implying their importance in folding and stabilizing prestin protein. Thus, p.A100T and p.P119S prestin missense variants that change the polarity of these residues may be detrimental to prestin function. Previously, we heterologously expressed p.A100T and p.P119S prestin in HEK293T cells and measured nonlinear capacitance (NLC), an electrical signature of electromotility (19, 20), to assess the functional impact of these variants. We found that both variants exhibited NLC, albeit with significantly reduced magnitudes compared to WT prestin (18). Prompted by these observations, we generated knock-in (KI) FVB/NJ mouse lines expressing p.A100T (*Slc26a5*^*+/A100T*^, *Slc26a5*^*A100T/A100T*^) and P119S prestin (*Slc26a5*^*+/P119S*^, *Slc26a5* ^*P119S/P119S*^) for in-depth functional characterization. We also obtained *Slc26a5*^*A100T/P119S*^ mice by crossing p.A100T and p.P119S prestin KI mice, as this compound heterozygosity was found in a human patient with congenital hearing loss (16).

To determine the physiological consequences of p.A100T and p.P119S prestin variants, we measured distortion product otoacoustic emissions (DPOAEs) that reflect intermodulation distortion associated with the OHC transducer channel. DPOAE measurements can detect subtle OHC dysfunction that may not yet be sufficient to manifest as hearing threshold shifts (21-25). Hence, DPOAEs are well suited for scrutinizing the pathogenic impact of prestin variants on OHC function *in vivo*. Briefly, DPOAEs are generated by the interaction of mechanical cochlear responses elicited by two pure tones, f_1_ and f_2_ (f_1_<f_2_). Compared to the basilar membrane traveling wave (propagation of sound-induced oscillations along the length of the cochlea) induced by f_2_, that for f_1_ peaks further toward the apex because of its lower frequency. DPOAEs are thought to be generated in the region of overlap of the two traveling waves produced by simultaneous presentation of the two stimulus tones. We measured DPOAE amplitude at 2f_1_-f_2_ with the f_2_/f_1_ ratio fixed at 1.2 to maximize this cubic distortion tone for sensitively detecting OHC dysfunction (26). **Figure 1A** shows frequency-dependent DPOAEs at 70 dB sound pressure level (SPL, L_1_ = L_2_ = 70 dB SPL), recorded in various *Slc26a5* genotypes indicated in each panel. At ∼1 month of age, *Slc26a5*^*A100T/A100T*^, *Slc26a5*^*P119S/P119S*^, and *Slc26a5*^*A100T/P119S*^ mice (**Fig. 1A**, green) all exhibited vastly reduced DPOAEs compared to age-matched WT (*Slc26a5*^*+/+*^) littermates (**Fig. 1A**, black). In contrast, *Slc26a5*^*+/A100T*^ and *Slc26a5*^*+/P119S*^ heterozygous littermates exhibited WT-like DPOAEs (**Fig. 1A**, left and middle panels, blue), recapitulating the recessive inheritance of DFNB61 hearing loss. At weaning (P18-22), relatively larger DPOAEs were observed in *Slc26a5*^*A100T/A100T*^, *Slc26a5*^*P119S/P119S*^, and *Slc26a5*^*A100T/P119S*^ mice for f_2_ less than ∼25 kHz (**Fig. 1A**, magenta). However, these responses were still significantly smaller than those of WT mice (the *p* values obtained by one-way ANOVA followed by Tukey’s multiple comparison post-test are provided in **Table S1**). We also performed sound level-dependent DPOAE measurements at f_2_ = 8, 12, 16, 24, and 32 kHz (f_2_/f_1_ = 1.2, L_1_ = L_2_+10 dB SPL) (**Fig. 1B**). At ∼1 month of age, DPOAEs in *Slc26a5*^*A100T/A100T*^, *Slc26a5*^*P119S/P119S*^, and *Slc26a5*^*A100T/P119S*^ mice (**Fig. 1B**, green) were smaller than in controls and sometimes statistically indistinguishable from coupler values (**Fig. 1B**, gray shades). Again, *Slc26a5*^*+/A100T*^ and *Slc26a5*^*+/P119S*^ heterozygotes (**Fig. 1B**, blue) were WT-like (**Fig. 1B**, black). At weaning, *Slc26a5*^*A100T/A100T*^, *Slc26a5*^*P119S/P119S*^, and *Slc26a5*^*A100T/P119S*^ mice showed detectable DPOAEs below 32 kHz (**Fig. 1B**, magenta); however, these responses were smaller compared to those seen in WT and heterozygotes (the *p* values obtained by one-way ANOVA followed by Tukey’s multiple comparison post-test are provided in **Table S2**).

**Figure 1.**
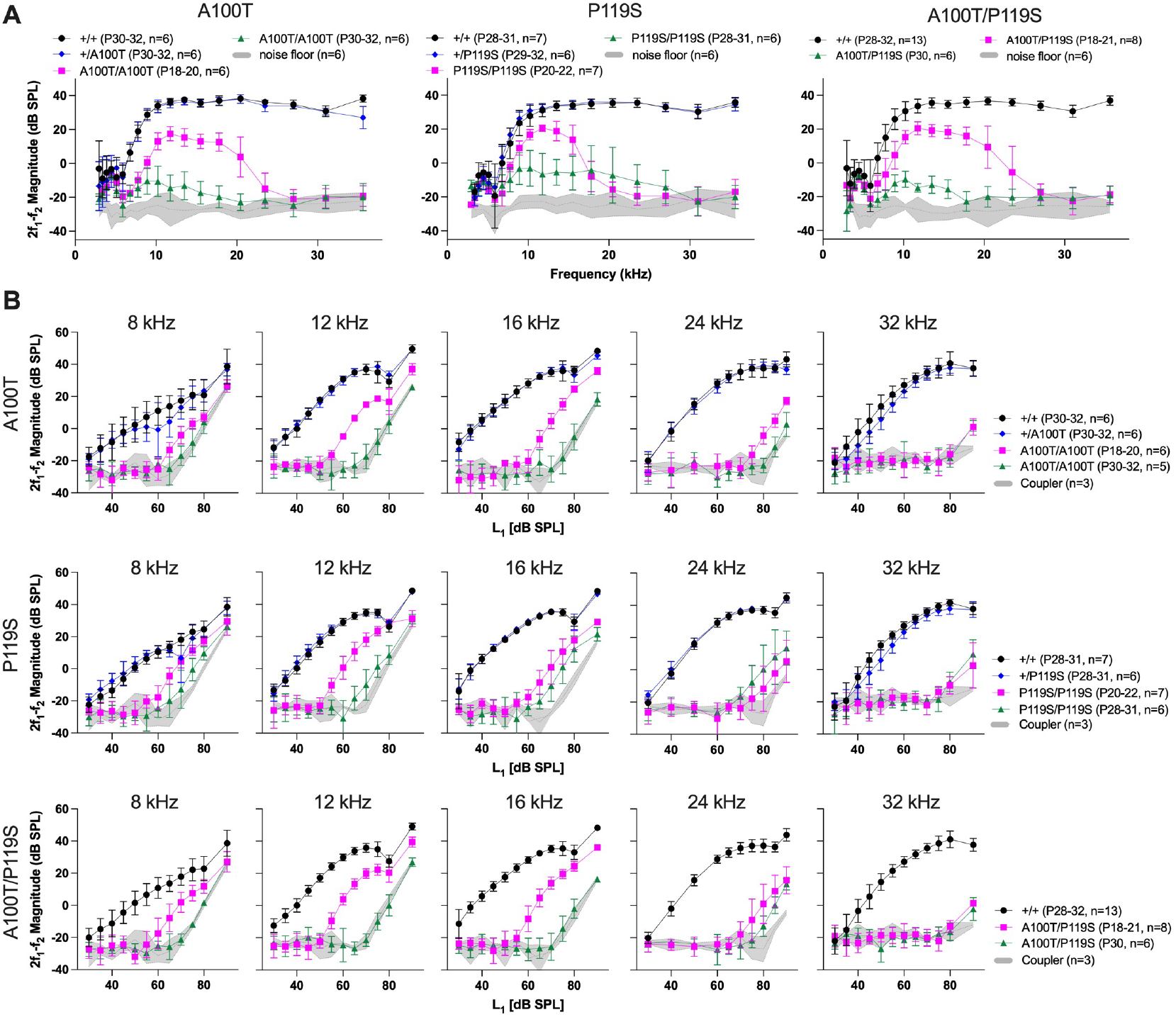
OHC function assessed using DPOAEs. (**A**) DPOAE iso-input functions. DPOAEs at 2f_1_-f_2_ in *Slc26a5*^*A100T/A100T*^ (left), *Slc26a5*^*P119S/P119S*^ (middle), and *Slc26a5*^*A100T/P119S*^ (right) mice, measured at weaning (P18-22, magenta) and at ∼1 month of age (P28-32, green), were compared to those of their WT (+/+, black) and heterozygous (A100T/+ or P119S/+, blue) littermates (P28-32). The f_2_/f_1_ ratio was fixed at 1.2, while the sound levels at f_1_ (L_1_) and f_2_ (L_2_) were set to 70 dB SPL. Error bars indicate SDs. Gray shading indicates the noise floor (mean ± SD). One-way ANOVA followed by Tukey’s multiple comparison post-test was performed at each frequency to obtain adjusted *p* values (**Table S1**). (**B**) DPOAE input-output functions. DPOAEs at 2f_1_-f_2_ were measured at various L_1_ input sound levels (L_2_=L_1_-10 dB SPL) at five different f_2_ frequencies (the f_2_/f_1_ ratio fixed at 1.2) in *Slc26a5*^*A100T/A100T*^ (top), *Slc26a5*^*P119S/P119S*^ (middle), and *Slc26a5*^*A100T/P119S*^ (bottom) at weaning (P18-22, magenta) and ∼1 month of age (P28-32, green). Results were compared to those of their WT (+/+, black) and heterozygous (A100T/+ or P119S/+, blue) littermates (P28-32). Error bars indicate SDs. Gray shading indicates coupler responses (mean ± SD), i.e., distortion in the sound. One-way ANOVA followed by Tukey’s multiple comparison post-test was performed at each L_1_ sound intensity to obtain adjusted *p* values (**Table S2**).

Using input-output functions shown in **Fig. 1B**, we defined DPOAE threshold as the level of f_1_ (L_1_) that produced a 2f_1_-f_2_ distortion product of 0 dB SPL. We then plotted the differences in the DPOAE thresholds as compared to those of WT (**Fig. 2A**) and compared them with the threshold shifts in the auditory brainstem responses (ABRs) (**Fig. 2B**). Both showed qualitatively similar frequency-dependent increases; however, it is notable that *Slc26a5*^*A100T/A100T*^, *Slc26a5*^*P119S/P119S*^, and *Slc26a5*^*A100T/P119S*^ mice all showed near-normal ABR thresholds at 8 kHz at weaning unlike the ∼20 dB threshold shifts found in DPOAEs at f_2_ of 8 kHz. This discrepancy is likely ascribed to the mechanism of DPOAE generation that is a nonlinear response generated by the interaction of f_1_ and f_2_ tones in the region of overlap at and basal to the f_2_ place (see above). In contrast, ABR responses were elicited by a single pure tone and do not depend on the production of intermodulation distortion products. The significantly elevated ABR thresholds at 12 kHz (**Fig. 2B**, magenta) suggest that OHCs with best frequencies greater than 8 kHz are impaired and thus could account for the reduced DPOAEs at f_2_ of 8 kHz. It should also be noted that the definition of DPOAE threshold (see above) differs from that for the ABRs.

**Figure 2.**
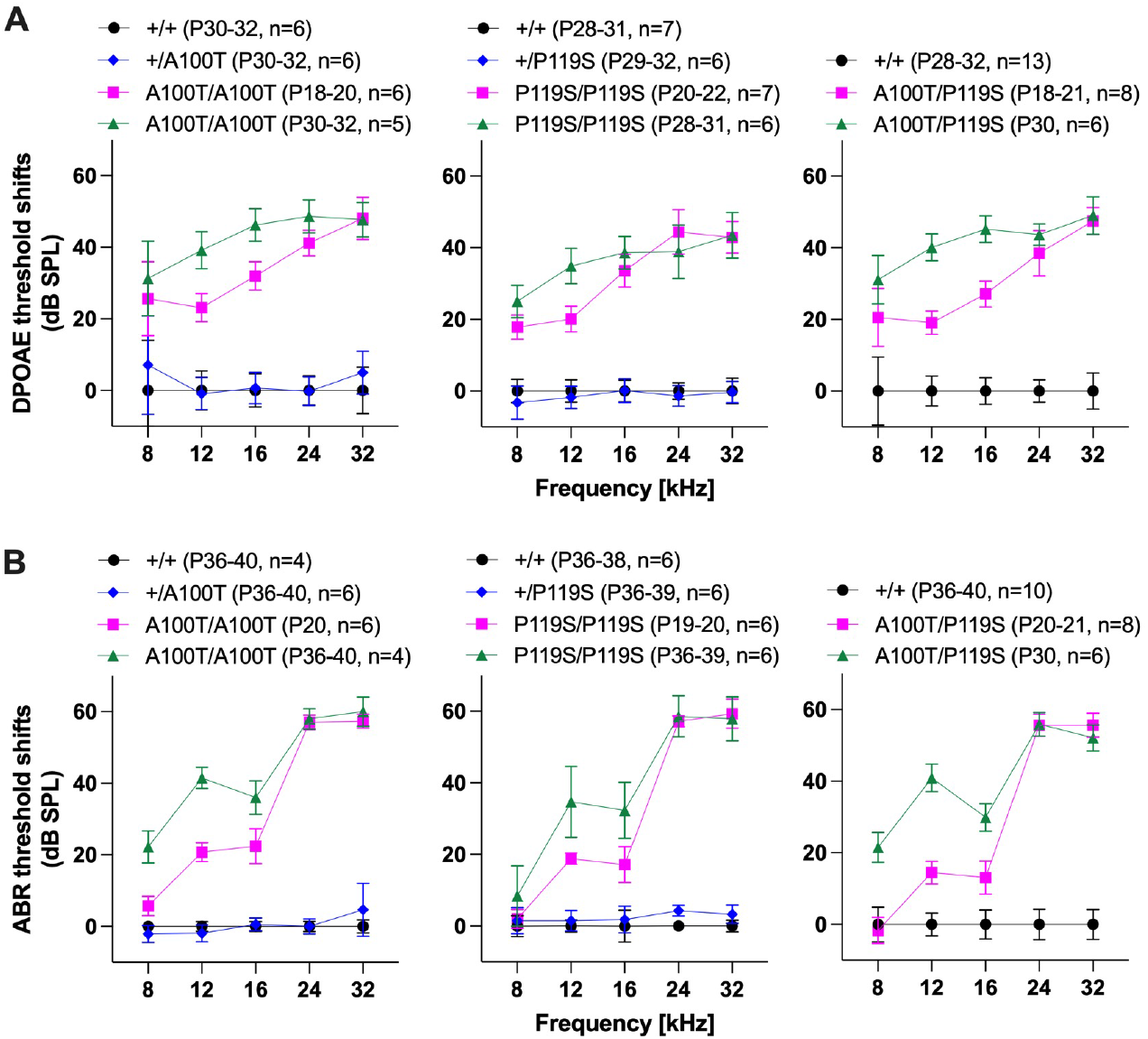
DPOAE and ABR threshold shifts. (**A**) DPOAE threshold shifts. L_1_ sound levels that produced a DPOAE at 2f_1_-f_2_ of 0 dB SPL (**Fig. 1B**) were defined as DPOAE thresholds, and their differences from WT controls were computed as threshold shifts and plotted with the same color code used in **Fig. 1**. (**B**) ABR threshold shifts. The differences in ABR thresholds (as compared to WT) were computed and plotted with the same color code as in panel A. In both panels, error bars indicate propagated errors calculated from SDs of the DPOAE and ABR threshold data. One-way ANOVA followed by Tukey’s multiple comparison post-test was performed at each frequency to obtain adjusted *p* values (**Tables S3 and S4** for panels A and B, respectively).

Taken together, these *in vivo* measurements demonstrate that both p.A100T and p.P119S induce progressive and frequency-dependent hearing loss, demonstrating their pathogenic contributions to congenital hearing loss found in human patients.

### Progressive OHC death contributes to the hearing loss in *Slc26a5*^*A100T/A100T*^, *Slc26a5*^*P119S/P119S*^, and *Slc26a5*_*A100T/P119S*_ mice

Premature OHC loss was found in mice lacking prestin or expressing dysfunctional prestin proteins (7, 8, 27, 28). Thus, we constructed cytocochleograms in order to learn whether OHC loss underlies the hearing deficit found in *Slc26a5*^*A100T/A100T*^, *Slc26a5*^*P119S/P119S*^, and *Slc26a5*^*A100T/P119S*^ mice (**Fig. 3**). At weaning, the OHC loss was minimal in homozygous and compound heterozygous mice (**Figs. 3B-3D**, magenta). By P30, however, *Slc26a5*^*A100T/A100T*^ and *Slc26a5*^*A100T/P119S*^ exhibited OHC loss in the basal region while retaining OHCs in the apical half of the cochlea (**Figs. 3B and 3D**, green). In *Slc26a5*^*P119S/P119S*^ homozygotes, OHC loss was more widespread at ∼1 month of age and progressed further apically at ∼6 weeks (**Fig. 3C**, green and brown, respectively). These observations indicate that p.A100T and p.P119S prestin variants have dual pathogenic roles: they primarily impair OHC function and secondarily induce OHC death.

**Figure 3.**
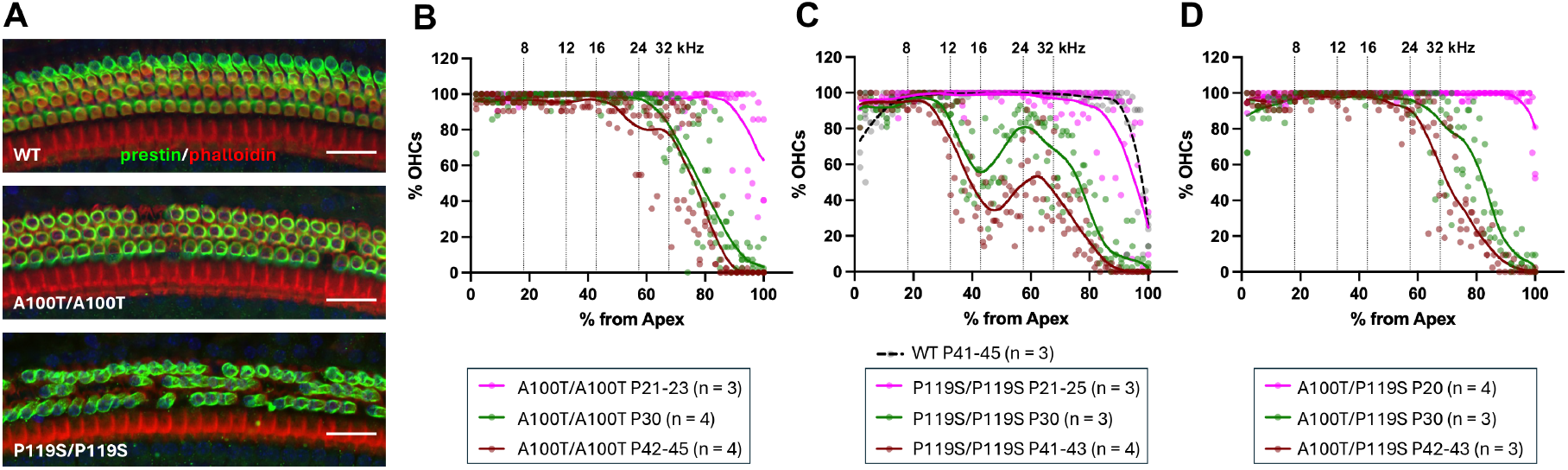
Progressive OHC loss in *Slc26a5*^*A100T/A100T*^, *Slc26a5*^*P119S/P119S*^, and *Slc26a5*^*A100T/P119S*^ mice. (**A**) Representative immunofluorescence microscopic images of the organ of Corti whole-mounts collected at P42. Images were taken from the middle turns at a distance ∼50% from the apex. Green, prestin; red, actin. Scale bars, 25 μm. OHC preservation was quantified at various postnatal ages for *Slc26a5*^*A100T/A100T*^ (**B**), *Slc26a5*^*P119S/P119S*^ (**C**), and *Slc26a5*^*A100T/P119S*^ (**D**). LOWESS fits (solid lines) are included to facilitate visualization of the extent of OHC loss in each graph. Cochlea locations corresponding to the frequencies tested for DPOAE and ABR measurements are also indicated. WT data at ∼6 weeks of age (P41-45) are included in panel C (black broken line).

### The p.A100T and p.P119S missense mutations severely reduce membrane expression of prestin protein but not its fast motor action

OHC loss does not account for the frequency-dependent DPOAE threshold shifts found in *Slc26a5*^*A100T/A100T*^, *Slc26a5*^*P119S/P119S*^, and *Slc26a5*^*A100T/P119S*^ mice at weaning and ∼1 month of age for f_2_ ≤12 kHz, which are at least 20 dB SPL. To further define the pathogenic roles of p.A100T and p.P119S prestin variants, OHCs isolated from p.A100T and p.P119S prestin KI mice at ∼1 month of age were inspected visually and electrophysiologically (**Fig. 4**). We found that p.A100T and p.P119S prestin variants do not alter the overall OHC appearance but significantly shorten cell length (**Figs. 4A and 4B**). Because prestin expression in the lateral membrane contributes to OHC length (7, 27), the shortening of OHCs indicates reduced membrane expression of p.A100T and p.P119S prestin variant proteins compared to WT. Whole-cell recordings in OHCs isolated from *Slc26a5*^*A100T/A100T*^, *Slc26a5*^*P119S/P119S*^, and *Slc26a5*^*A100T/P119S*^ mice (**Figs. 4C and 4D**) revealed large reductions in electromotility (**Fig. 4E**), C_lin_ that correlates with cell size (**Fig. 4F**), and the magnitude of NLC (NLC_pk_, **Fig. 4F**). Although statistically significant, the reduction in the voltage sensitivity (α) found in OHCs isolated from *Slc26a5*^*A100T/P119S*^ mice was small. Small but statistically significant changes were also found in the voltage operating point (V_pk_) in OHCs isolated from *Slc26a5*^*A100T/A100T*^, *Slc26a5*^*+/P119S*^, *Slc26*

**Figure 4.**
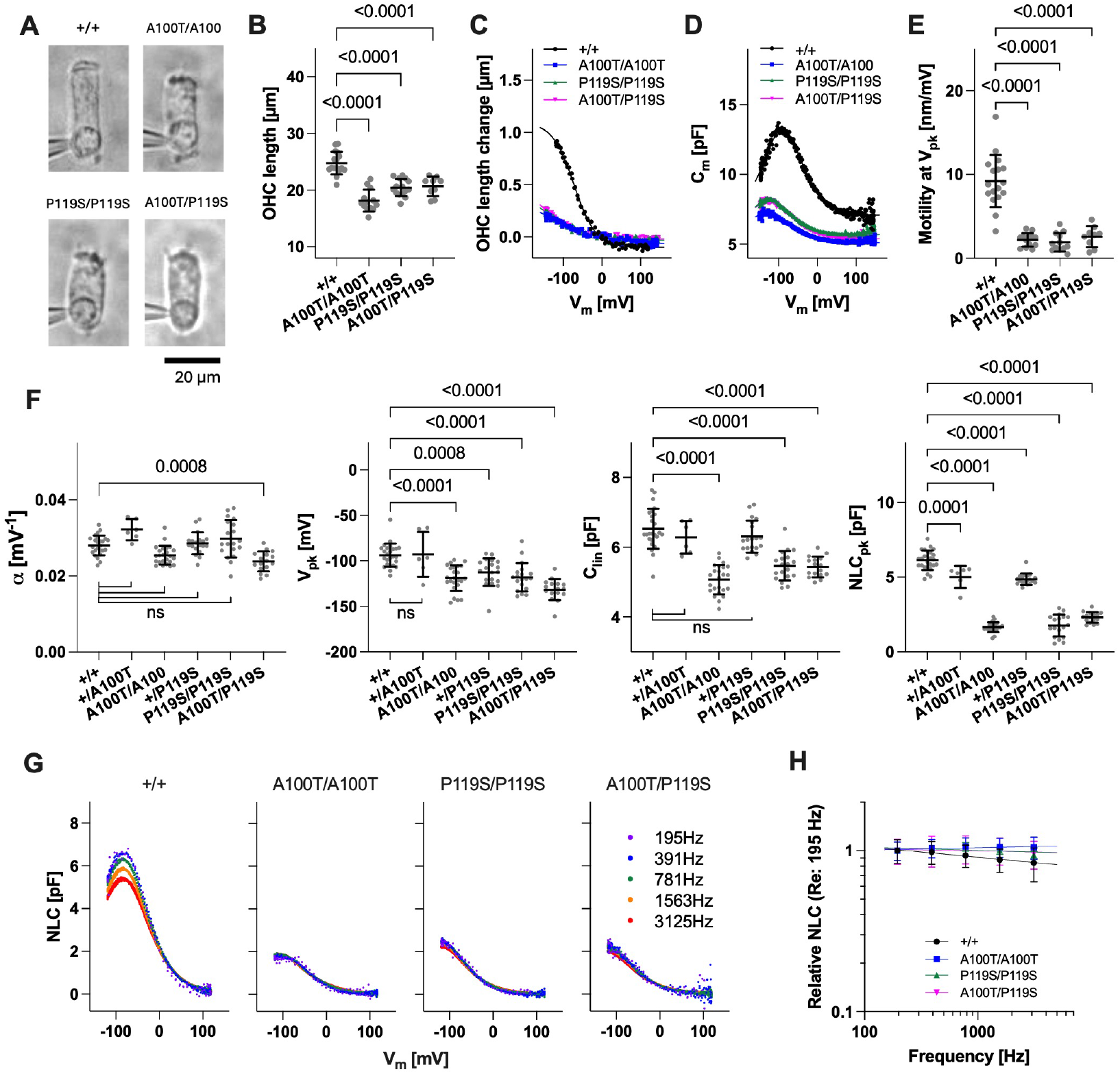
Electromotility and NLC recordings in OHCs. OHCs were isolated from the apical region (corresponding to 4 to 10 kHz) of the cochlea of *Slc26a5*^*+/+*^, *Slc26a5*^*A100T/A100T*^, *Slc26a5*^*P119S/P119S*^, and *Slc26a5*^*A100T/P119S*^ mice at P28-37 for electromotility and NLC recordings in the whole-cell configuration. (**A**) Representative microscopic images of isolated OHCs. Portions of whole-cell recording glass pipettes are also seen at the basal regions of OHCs. (**B**) A summary of OHC length. (**C, D**) Examples of electromotility and NLC recordings. Solid lines indicate two-state Boltzmann curve fits. Summaries of electromotility (**E**) and NLC parameters (**F**). α is the slope factor. V_pk_ is the voltage at which the maximum NLC (NLC_pk_) is attained. C_lin_ is the linear capacitance that correlates with cell size. The *p* values were obtained by one-way ANOVA followed by Tukey’s multiple comparison post-test. ns, *p* ≥ 0.05. Error bars indicate SDs. (**G**) Examples of stimulus frequency-dependent NLC measured at five different f_1_ frequencies indicated with the corresponding f_2_ frequencies being twice as large as f_1_ (i.e., f_2_=390, 780, 1563, 3125, 6250 Hz). (**H**) Stimulus frequency-dependent relative NLC_pk_ (normalized to the magnitudes of NLC_pk_ at 195 Hz. Error bars indicate propagated errors calculated from SDs of the NLC_pk_ data before normalization.

*a5*^*P119S/P119S*^, and *Slc26a5*^*A100T/P119S*^ mice (**Fig. 4F**). However, it is unlikely that these hyperpolarizing V_pk_ shifts have any pathogenic impact because mice expressing prestin with ‘C1’ triple mutations (29) that negatively shift V_pk_ >50 mV retain normal hearing (15, 30). We also measured NLC in a stimulus frequency-dependent manner to examine the impact of p.A100T and p.P119S on prestin’s motor speed (**Figs. 4G and 4H**). As evident in these figures, we did not find any signs of reduced motor kinetics in OHCs isolated from *Slc26a5*^*A100T/A100T*^, *Slc26a5*^*P119S/P119S*^, and *Slc26a5*^*A100T/P119S*^ mice (*p* > 0.05, *F*-test), suggesting that reduced membrane expression of p.A100T and p.P119S prestin proteins underlies hearing loss in *Slc26a5*^*A100T/A100T*^, *Slc26a5*^*P119S/P119S*^, *Slc26a5*^*A100T/P119S*^ mice at weaning and ∼1 month of age for f_2_ ≤12 kHz.

To uncover the reason why DPOAE and ABR threshold shifts were smaller at weaning compared to ∼1 month of age for the apical region (≤ 12 kHz) where OHC loss is minimal at both ages (**Fig. 3**), we repeated whole-cell recordings in OHCs isolated from *Slc26a5*^*A100T/A100T*^ and *Slc26a5*^*P119S/P119S*^ mice at weaning and compared the results with those obtained at ∼1 month of age (**Fig. 5**). As anticipated, differences in the voltage sensitivity (α) and operating point (V_pk_) were not detected or very small (**Fig. 5A**). However, we found larger C_lin_ values at weaning (**Fig. 5A**). This observation corroborates with direct visual inspection (i.e., longer OHCs at weaning, **Fig. 5B**), suggesting a greater cell membrane expression of p.A100T and p.P119S prestin protein at the younger age. Although variable, the magnitudes of NLC_pk_ and electromotility tended to be greater at weaning (**Figs. 5A and 5C**).

**Figure 5.**
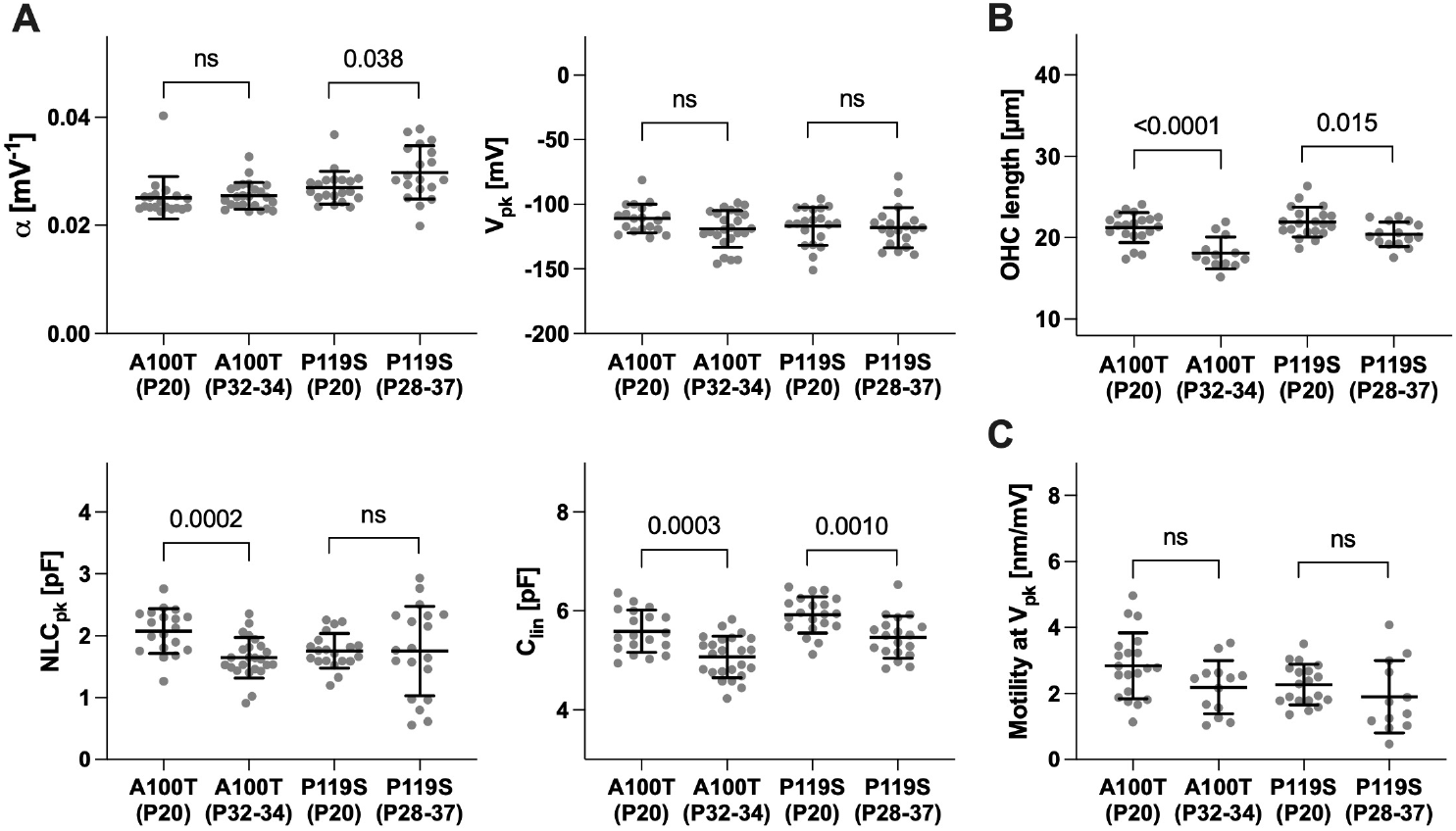
Reduction of OHC function after weaning. Summaries of NLC parameters (**A**), OHC length (**B**), and electromotility (**C**) measured in OHCs isolated from *Slc26a5*^*A100T/A100T*^ and *Slc26a5*^*P119S/P119S*^ homozygous mice at weaning (P20) and ∼1 month of age. Error bars indicate SDs. A two-tailed t-test was performed to assess the significance of the difference between the mean values. ns, *p* ≥ 0.05.

Collectively, these observations indicate that hearing loss in *Slc26a5*^*A100T/A100T*^, *Slc26a5*^*P119S/P119S*^, and *Slc26a5*^*A100T/P119S*^ mice is primarily ascribed to progressively reducing membrane expression of p.A100T and p.P119S prestin proteins and not to altered motor characteristics.

### The minimum prestin motor activity required for normal hearing sensitivity

Compared to WT, the magnitudes of OHC electromotility at weaning and ∼1 month of age were 31% and 24% for *Slc26a5*^*A100T/A100T*^ mice and 25% and 21% for *Slc26a5*^*P119S/P119S*^ mice, respectively (**Figs. 4E and 5C**). This means that near-normal hearing sensitivity can be attained with ∼30% OHC electromotility at the lowest frequency tested in the present study, 8 kHz (**Fig. 2B**). Our whole-cell recordings were limited to OHCs isolated from the apical region corresponding to 4-10 kHz due to the difficulty in isolating OHCs from higher frequency regions. However, if % reductions of OHC electromotility were to be similar at the other cochlear locations, our results would suggest that the minimum prestin motor activity required for attaining a normal hearing threshold would be greater at higher frequencies. This view is in line with previous studies showing that 44% of OHC electromotility supports normal hearing thresholds at 4-22 kHz but not at 32 kHz (13) and that 54% of OHC electromotility is sufficient to maintain normal cochlear amplification at all tested frequencies from 2 to 45 kHz (14, 15). These observations suggest that DFNB61 hearing loss could be ameliorated without restoring OHC electromotility to the WT level.

### The pathogenic role of the p.A100T and p.P119S prestin variants

Chloride is an extrinsic cofactor essential for prestin’s motor function (29, 31, 32). The intracellular chloride concentration, [Cl^-^]_i_, in mature OHCs is estimated to be 10 mM or less (33). It is conceivable that [Cl^-^]_i_ is higher in early postnatal OHCs because previous transcriptomics studies revealed postnatal decline of *Slc12a2* (NKCC1, Cl^-^ uptake) and increase of *Slc12a7* (KCC4, Cl^-^ exclusion) (34, 35). We speculate that p.A100T and p.P119S impair protein stability, resulting in reduced cell membrane targeting of the otherwise functional prestin protein, and that the binding of chloride may stabilize the p.A100T and p.P119S variants augmenting their membrane expression in young mice. To explore this possibility, we expressed p.A100T and p.P119S prestin in HEK293T cells with and without a second missense mutation, p.S396D, that mimics chloride binding (12), and measured NLC (**Fig. 6**). Consistent with a recent report (18), heterologous expression of p.A100T and p.P119S prestin conferred very small NLC on HEK293T cells compared to WT prestin (**Fig. 6**). Introduction of p.S396D increased the magnitude of NLC (**Fig. 6**), implying that mimicking chloride binding facilitated membrane targeting of p.A100T and p.P119S prestin. We also performed similar experiments using p.A100T and p.P119S prestin-expressing HEK293T cells that were cultured in the presence of 10 mM of the prestin inhibitor, salicylate, that competes for the chloride binding site in prestin. Salicylate was removed before NLC recordings. This pharmacological pretreatment was also found effective in significantly augmenting NLC (**Fig. 6**).

**Figure 6.**
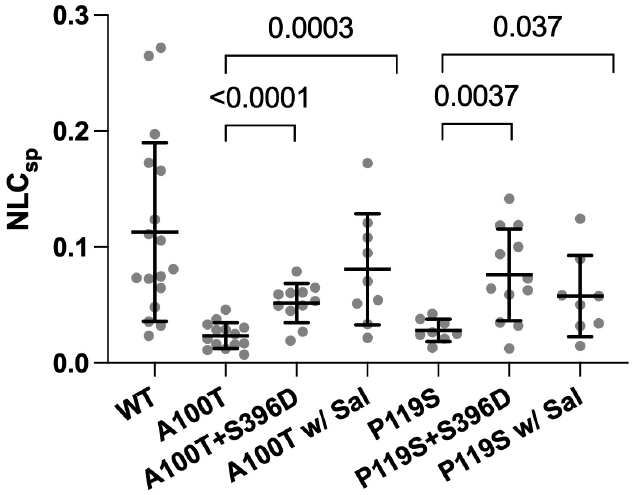
Genetic and pharmacological augmentation of cell membrane targeting of p.A100T and p.P119S prestin. Expression of p.A100T and p.119S prestin was induced in stable HEK293T cell lines using 1 μg/mL doxycycline with or without 10 mM salicylate for 2 days. NLC was measured after washing the cells with a bath solution lacking salicylate. Error bars indicate SDs. Improvement in NLC was assessed by a two-tailed t-test.

These observations indicate that the progressive DPOAE/ABR threshold shifts at ≤12 kHz (**Fig. 2**) may relate to the reduction in membrane expression of prestin protein (**Fig. 5**) associated with the presumptive postnatal gradual reduction of [Cl^-^]_i_. The significant improvement of NLC in HEK293T cells pretreated with salicylate supports the potential for pharmacological restoration of hearing in DFNB61 patients.

## DISCUSSION

The present study defined the functional consequences of p.A100T and p.P119S prestin variants using KI mouse models. We found that homozygous and compound heterozygous mice expressing these prestin variants suffer from congenital and progressive hearing loss, demonstrating that p.A100T and p.P119S prestin variants are indeed pathogenic. We also found that heterozygous p.A100T and p.P119S KI mice have normal hearing sensitivity across frequency, recapitulating the recessive inheritance of DFNB61 hearing loss. Although the p.A100T and p.P119S missense changes only slightly altered the voltage sensitivity and operating point, they vastly reduced membrane expression, implying that these variants may impair the stability of prestin protein leading to rapid degradation. Interestingly, the NLC of p.A100T and p.P119S prestin variants was *less* frequency dependent (i.e., had faster motor speed) compared to WT. Although the differences were not statistically significant, it is tempting to speculate that destabilization of prestin protein by p.A100T and p.P119S (i.e., an increase in the structural free energy) reduces the energy barrier (activation energy) between prestin’s contracted and expanded states, resulting in faster motor kinetics. Hence, the signs of increased motor kinetics are consistent with the suggestion that p.A100T and p.P119S destabilize prestin protein.

One of the prominent findings was that a normal ABR threshold at 8 kHz, the lowest frequency tested in the present study, was attained with vastly reduced OHC electromotility as low as 25-31% of WT. Elevated thresholds at higher frequencies (12, 16, 24, and 32 kHz) suggest two possibilities: either larger electromotility or cochlear gain is required at higher frequencies, or the reduction in electromotility (%WT) is greater in higher frequency regions. We speculate that the former may be the case because it seems unlikely that the membrane targeting efficiencies of p.A100T and p.P119S prestin proteins differ significantly between OHCs at different locations along the cochlear partition. Given the low-pass nature of prestin motor activity (36), the tolerance of cochlear amplification to reductions in prestin’s motor activity may be smaller in basal cochlear regions. This speculation is in line with previous studies showing that 44% of OHC electromotility is sufficient to retain normal hearing thresholds at 4-22 kHz but not at 32 kHz (13) and that 54% of OHC electromotility is sufficient for normal cochlear amplification at 2-45 kHz (14, 15). The large tolerance of cochlear function to reduced prestin motor activity may account for the scarcity of pathogenic *SLC26A5* variants identified in human patients.

Collectively, these results point to the possibility of ameliorating DFNB61 hearing loss by partially restoring OHC electromotility. For example, the significant increases of NLC in p.A100T and p.P119S prestin-expressing HEK293T cells pretreated with salicylate (**Fig. 6**) provide an exciting possibility for pharmacological restoration of hearing in patients harboring these pathogenic prestin variants. Although salicylate *per se* will not be a practical choice for therapeutics as it inhibits prestin activity, identification of small molecules that enhance prestin protein stability and/or membrane targeting could potentially restore pathogenic effects of prestin variants such as p.A100T and p.P119S.

Considering the large threshold shifts documented in mice lacking prestin, it is counterintuitive that ∼60% reduction in prestin-mediated OHC electromotility barely or only slightly affects hearing sensitivity across frequency. To address this enigma, we recently proposed an OHC electromotility mechanism powered by intracellular turgor pressure in which prestin acts as a voltage-dependent stiffness modulator. In this scenario, the prestin-associated axial stiffness is largely insensitive to large reductions of prestin (15). The cylindrical shape of OHCs whose lateral membranes are not physically in contact with other structures is ideal for this hypothetical turgor pressure-powered electromotility mechanism to work efficiently. In addition, the magnitude of OHC electromotility can be readily increased or decreased by changing turgor pressure, which is consistent with previous studies experimentally demonstrating the dependence of OHC electromotility on turgor (37, 38). It has also been shown that perfusion of hypotonic perilymph, which is expected to increase turgor pressure, enhances cochlear amplification *in vivo* (39). We speculate that increased turgor pressure may be important at higher frequency locations to maintain sufficient OHC electromotility that is otherwise attenuated due to the intrinsic low-pass nature of prestin motor activity (36). It is also possible that increased turgor contributes to the vulnerability of basal OHCs to mechanical insults. Premature OHC loss, beginning at the base and progressing apically, was reported in mice lacking prestin (7, 8), in mice expressing a virtually nonelectromotile p.V499G/p.Y501H prestin mutation (10, 27), and in the p.R130S prestin variant that slows OHC electromotility (12, 28). The present study also found that mice expressing p.A100T and p.P119S prestin variants suffer from premature OHC loss. It is conceivable that turgor pressure may be overly upregulated in OHCs with loss of or with reduced prestin function, increasing susceptibility to sound-elicited mechanical stress.

Finally, a recent study showed that the loss of TMEM63B, a stretch-activated nonselective cation channel, induces OHC loss and results in a progressive hearing loss in mice (40). This report suggests that TMEM63B regulates turgor pressure, and that turgor is dynamically regulated to optimize the OHC’s mechanical contribution to cochlear gain. In addition, there are other membrane channels and transporters that are expressed in mature OHCs (Gene Expression Omnibus: GSE111349) and are known to regulate cell volume (41), including those responsible for cell volume increase (e.g., Na^+^/H^+^ exchangers, Cl^-^/HCO_3_^-^ exchangers, and Na^+^/K^+^/2Cl^-^ cotransporters, etc.) and cell volume decrease (e.g., K^+^/Cl^-^ cotransporters, volume-regulated anion channels, Ca^2+^-activated Cl^-^ channels, Ca^2+^-activated K^+^ channels, and voltage-independent K^+^ channels, etc.). Further characterization of the p.A100T and p.P119S mouse models may help to identify key molecular players responsible for OHC turgor regulation and for validating the proposal that turgor regulation modulates OHC electromotility *in vivo*.

## MATERIALS AND METHODS

### Animals

Prestin mouse models harboring p.A100T or p.P119S were generated on the FVB/NJ background using CRISPR/Cas9 at Northwestern University’s Transgenic and Targeted Mutagenesis Laboratory. The mouse lines were maintained by heterozygous mating and genotyped by Transnetyx (Cordova, TN). Compound heterozygous mice for p.A100T and p.P119S (*Slc26a5*^*A100T/P119S*^) were obtained by crossing homozygous p.A100T and p.P119S mutants. All procedures and protocols were approved by Northwestern University’s Institutional Animal Care and Use Committee (IS00005343) and by NIDCD.

### Hearing tests

Anesthetized (100 mg/Kg ketamine/10 mg/Kg xylazine IP) animals were screened at various ages to determine the hearing phenotype using noninvasive methods as in a previous study (12). Both male and female mice were tested, and all recordings were from the animal’s left ear in a sound-isolated and electrically shielded chamber. A heating pad was used to maintain body temperature at approximately 37°C. Distortion product otoacoustic emissions (DPOAEs) were recorded using a custom probe placed close to the eardrum (42). DPOAE input-output functions were collected at various f_2_ frequencies (f_2_/f_1_=1.2), where L_1_ = L_2_ + 10 dB SPL, and used to determine DPOAE thresholds defined as the level of f_1_ that produced a 2f_1_-f_2_ component of 0 dB SPL. Iso-input functions collected at L_1_=L_2_=70 dB SPL were also acquired with a fixed frequency ratio (f_2_/f_1_=1.2) and with f_2_ increasing from 4-36 kHz. Auditory brainstem responses (ABRs), which reflect the integrity of the auditory nerve and brainstem pathways, were recorded using three stainless steel needle electrodes inserted at the vertex, the mastoid, and the contralateral ear. ABRs were elicited using 5 msec tone bursts including the 0.5 msec rise/fall times and presented with a repetition rate of 20/sec. Responses were averaged (512 samples), amplified and filtered (300-2,000 Hz). Thresholds were obtained at various frequencies (8-32 kHz) by noting the stimulus level at which the waveform disappeared into the noise floor. Data are provided as threshold differences between mutants and controls.

### Cytocochleograms

Mice were cardiac perfused with 4% paraformaldehyde and cochleae extracted. After post-fixation and decalcification, cochleae were dissected following the Eaton-Peabody Laboratory cochlear dissection protocol for whole mounts (43). Cochleae were stained with anti-prestin antisera (44) followed by goat anti-rabbit Alexa Fluor 488 secondary antibody (ThermoFisher), together with Alexa Fluor 568 phalloidin and Hoechst 33342 (ThermoFisher) for actin and nuclei, respectively. Images were taken using a Leica DM IRB fluorescence microscope or Keyence BZ-X800 with 10X and 20X objectives for cytocochleograms, as well as with Nikon A1R+ laser scanning confocal microscope as previously described (12). Cochlear locations corresponding to the frequencies tested for the DPOAE and ABR measurements were determined based on the Müller et al. frequency place map (45). LOWESS smooth fits were included in summary figures to facilitate visual inspection of the results.

### Generation of stable cell lines

Mouse prestin constructs (wildtype, p.A100T, p.P119S, p.A100T+p.S396D, p.P119S +p.S396D) with a C-terminally attached mTurquoise2 (mTq2) were cloned into a pSBtet-pur vector, and HEK293T-based stable cell lines that express these prestin constructs in a doxycycline-dependent manner were established as in previous studies (18, 28, 46-50). Expression of the prestin constructs was induced by 1 μg/mL doxycycline with or without 10 mM salicylate added to the culture media two days before experiments.

### NLC and electromotility measurements

OHCs were isolated as described previously (14). Whole-cell recordings were performed at room temperature using the Axopatch 200B amplifier (Molecular Devices) with a 10 kHz low-pass filter. Recording pipettes pulled from borosilicate glass were filled with a solution containing (mM): 140 CsCl, 2 MgCl_2_, 10 EGTA, and 10 HEPES (pH 7.4). Cells were bathed in an extracellular solution containing (mM): 120 NaCl, 20 TEA-Cl, 2 CoCl_2_, 2 MgCl_2_, 10 HEPES (pH 7.4). Osmolality was adjusted to 309 mOsmol/kg with glucose. While recording, the intracellular pressure was not adjusted. The electric current response to a sinusoidal voltage stimulus (2.5 Hz, 120 or 150 mV amplitude) superimposed with two higher frequency stimuli (390.6 and 781.2 Hz, 10 mV amplitude) was recorded by jClamp (SciSoft Company). For stimulus frequency-dependent C_m_ measurements, f_1_ was set at 195.3 (f_2_=390.6), 390.6 (f_2_=781.3), 781.3 (f_2_=1562.5), 1562.5 (f_2_=3125.0), and 3125.0 (f_2_=6250.0) Hz (14, 51) and a fast Fourier transform-based admittance analysis was used to determine C_m_ (52). Since both f_1_ and f_2_ contribute to C_m_ measurement, f_1_ frequencies were used to report frequency-dependent NLC data. OHC electromotility was captured using a Prosilica GE680 camera (Allied Vision) and the recorded sequential images analyzed using ImageJ as described previously (53).

### NLC and electromotility data analysis

Voltage-dependent C_m_ data were analyzed using the following two-state Boltzmann equation (see Takahashi and Homma, 2024 (50) for derivation):

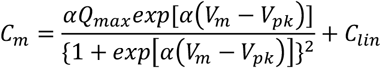

where α is the slope factor of the voltage-dependence of charge transfer, Q_max_ is the maximum charge transfer, V_m_ is the membrane potential, V_pk_ is the voltage at which the maximum charge movement is attained, and C_lin_ is the linear capacitance. Electromotility data were analyzed using the following equation:

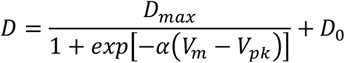

where D_max_ is the maximum cell length change, and D_0_ is a base reference point at which an OHC shows zero cell contraction (at infinitely hyperpolarized membrane potential).

### Calculations of propagated errors

The uncertainties (σ) associated with subtraction and division computations to determine the differences in DPOAE/ABR thresholds as compared to the wildtype controls and relative magnitudes of NLC measured at different stimulus frequencies with respect to those measured at 195 Hz (f_2_ = 391 Hz) were calculated by the following equations:

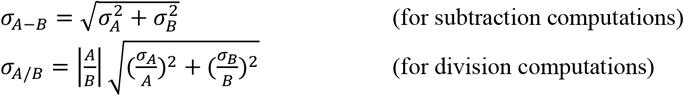

where A and B are the mean values with associated errors (standard deviations), σ_A_ and σ_B_, respectively.

### Statistical analyses

Statistical analyses were performed using Prism (GraphPad Software). The Student’s t-test was used for comparisons between two groups. One-way ANOVA combined with Tukey’s post hoc test was used for multiple comparisons. *F*-tests were performed to find differences in the stimulus frequency-dependence of NLC. In all statistical analyses, *p* < 0.05 was considered significant.

## Acknowledgments

This work was supported by an NIH grant DC017482 (to KH) and by the Hugh Knowles Center. Some of the imaging work was performed at the Northwestern University Center for Advanced Microscopy generously supported by NCI CCSG P30 CA060553 awarded to the Robert H Lurie Comprehensive Cancer Center.

## FIGURES AND LEGENDS

**Table S1.**
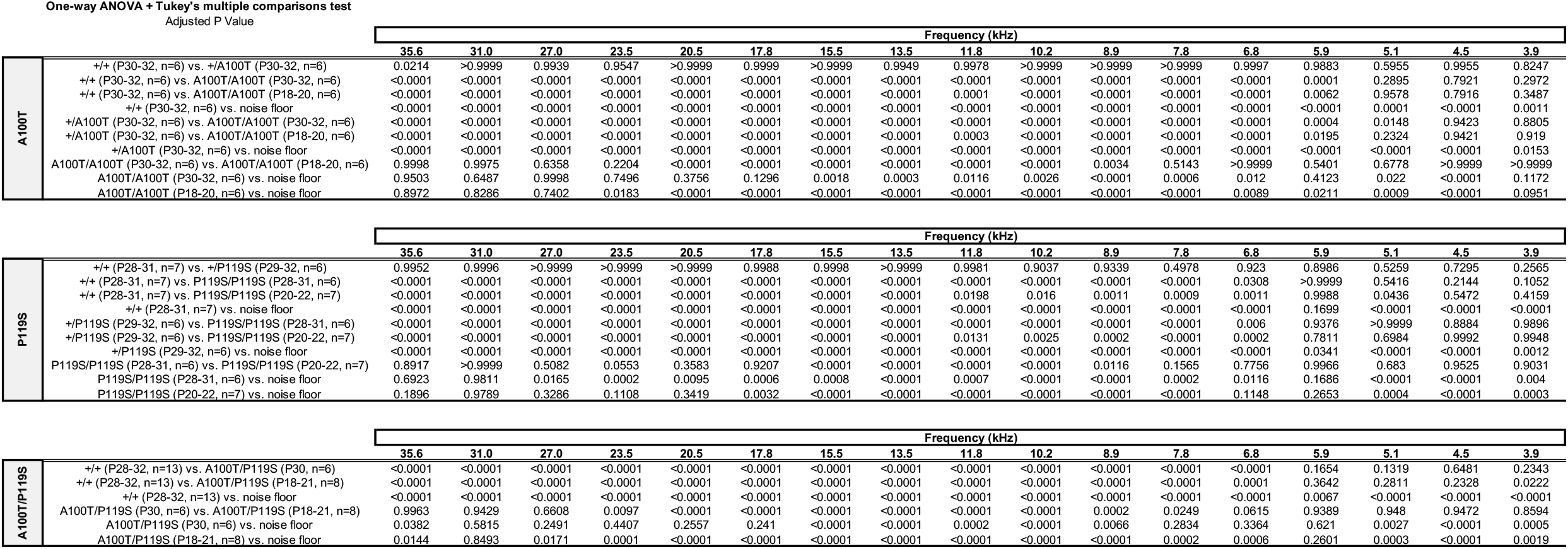

**Table S2.**
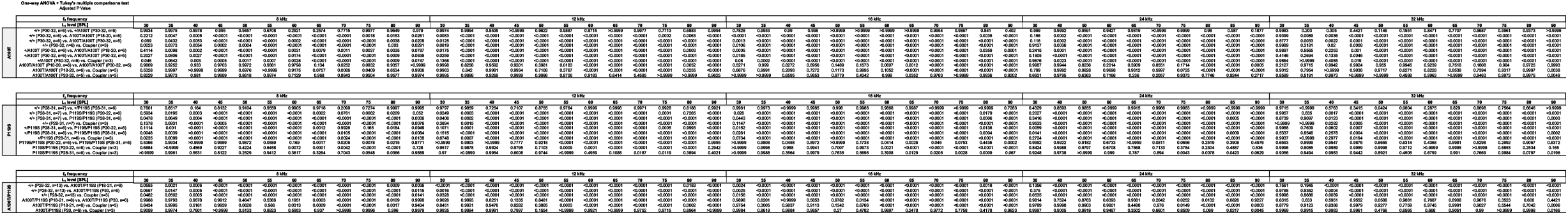

**Table S3.** One-way ANOVA + Tukey's multiple comparisons test Adjusted P Value

|  |  | f<sub>2</sub> frequency |  |  |  |  |
| --- | --- | --- | --- | --- | --- | --- |
|  |  | 8 kHz | 12 kHz | 16 kHz | 24 kHz | 32 kHz |
| A100T | +/+ (P30-32, n=6) vs. +/A100T (P30-32, n=6) | 0.428 | 0.9803 | 0.9924 | 0.9999 | 0.1771 |
|  | +/+ (P30-32, n=6) vs. A100T/A100T (P18-20, n=6) | <0.0001 | <0.0001 | <0.0001 | <0.0001 | <0.0001 |
|  | +/+ (P30-32, n=6) vs. A100T/A100T (P30-32, n=5) | <0.0001 | <0.0001 | <0.0001 | <0.0001 | <0.0001 |
|  | +/+ (P30-32, n=6) vs. Coupler (n=3) | <0.0001 | <0.0001 | <0.0001 | <0.0001 | <0.0001 |
|  | +/A100T (P30-32, n=6) vs. A100T/A100T (P18-20, n=6) | 0.0013 | <0.0001 | <0.0001 | <0.0001 | <0.0001 |
|  | +/A100T (P30-32, n=6) vs. A100T/A100T (P30-32, n=5) | 0.0001 | <0.0001 | <0.0001 | <0.0001 | <0.0001 |
|  | +/A100T (P30-32, n=6) vs. Coupler (n=3) | 0.0003 | <0.0001 | <0.0001 | <0.0001 | <0.0001 |
|  | A100T/A100T (P18-20, n=6) vs. A100T/A100T (P30-32, n=5) | 0.6739 | <0.0001 | <0.0001 | 0.0015 | 0.9998 |
|  | A100T/A100T (P18-20, n=6) vs. Coupler (n=3) | 0.603 | <0.0001 | <0.0001 | 0.0005 | 0.4473 |
|  | A100T/A100T (P30-32, n=5) vs. Coupler (n=3) | 0.9981 | >0.9999 | 0.9949 | 0.8071 | 0.4413 |
|  |  | 8 kHz | 12 kHz | 16 kHz | 24 kHz | 32 kHz |
| P119S | +/+ (P30-32, n=6) vs. +/A100T (P30-32, n=6) | 0.3614 | 0.8051 | >0.9999 | 0.984 | >0.9999 |
|  | +/+ (P30-32, n=6) vs. A100T/A100T (P18-20, n=6) | <0.0001 | <0.0001 | <0.0001 | <0.0001 | <0.0001 |
|  | +/+ (P30-32, n=6) vs. A100T/A100T (P30-32, n=5) | <0.0001 | <0.0001 | <0.0001 | <0.0001 | <0.0001 |
|  | +/+ (P30-32, n=6) vs. Coupler (n=3) | <0.0001 | <0.0001 | <0.0001 | <0.0001 | <0.0001 |
|  | +/A100T (P30-32, n=6) vs. A100T/A100T (P18-20, n=6) | <0.0001 | <0.0001 | <0.0001 | <0.0001 | <0.0001 |
|  | +/A100T (P30-32, n=6) vs. A100T/A100T (P30-32, n=5) | <0.0001 | <0.0001 | <0.0001 | <0.0001 | <0.0001 |
|  | +/A100T (P30-32, n=6) vs. Coupler (n=3) | <0.0001 | <0.0001 | <0.0001 | <0.0001 | <0.0001 |
|  | A100T/A100T (P18-20, n=6) vs. A100T/A100T (P30-32, n=5) | 0.003 | <0.0001 | 0.0604 | 0.2632 | 0.9989 |
|  | A100T/A100T (P18-20, n=6) vs. Coupler (n=3) | <0.0001 | <0.0001 | <0.0001 | 0.3705 | 0.0234 |
|  | A100T/A100T (P30-32, n=5) vs. Coupler (n=3) | 0.16 | 0.4097 | 0.0077 | 0.0134 | 0.0299 |
|  |  | 8 kHz | 12 kHz | 16 kHz | 24 kHz | 32 kHz |
| A100T/P119S | +/+ (P28-32, n=13) vs. A100T/P119S (P18-21, n=8) | <0.0001 | <0.0001 | <0.0001 | <0.0001 | <0.0001 |
|  | +/+ (P28-32, n=13) vs. A100T/P119S (P30, n=6) | <0.0001 | <0.0001 | <0.0001 | <0.0001 | <0.0001 |
|  | +/+ (P28-32, n=13) vs. Coupler (n=3) | <0.0001 | <0.0001 | <0.0001 | <0.0001 | <0.0001 |
|  | A100T/P119S (P18-21, n=8) vs. A100T/P119S (P30, n=6) | 0.0041 | <0.0001 | <0.0001 | 0.0768 | 0.8162 |
|  | A100T/P119S (P18-21, n=8) vs. Coupler (n=3) | 0.0209 | <0.0001 | <0.0001 | 0.0002 | 0.1277 |
|  | A100T/P119S (P30, n=6) vs. Coupler (n=3) | 0.9997 | 0.8977 | 0.7285 | 0.0642 | 0.4874 |

**Table S4.** One-way ANOVA + Tukey's multiple comparisons test Adjusted P Value

|  |  | frequency |  |  |  |  |
| --- | --- | --- | --- | --- | --- | --- |
|  |  | 8 kHz | 12 kHz | 16 kHz | 24 kHz | 32 kHz |
| A100T | +/+ (P36-40, n=4) vs. +/A100T (P36-40, n=6) | 0.632 | 0.5486 | 0.9934 | >0.9999 | 0.4024 |
|  | +/+ (P36-40, n=4) vs. A100T/A100T (P20, n=6) | 0.02 | <0.0001 | <0.0001 | <0.0001 | <0.0001 |
|  | +/+ (P36-40, n=4) vs. A100T/A100T (P36-40, n=4) | <0.0001 | <0.0001 | <0.0001 | <0.0001 | <0.0001 |
|  | +/A100T (P36-40, n=6) vs. A100T/A100T (P20, n=6) | 0.0006 | <0.0001 | <0.0001 | <0.0001 | <0.0001 |
|  | +/A100T (P36-40, n=6) vs. A100T/A100T (P36-40, n=4) | <0.0001 | <0.0001 | <0.0001 | <0.0001 | <0.0001 |
|  | A100T/A100T (P20, n=6) vs. A100T/A100T (P36-40, n=4) | <0.0001 | <0.0001 | <0.0001 | 0.8438 | 0.7954 |
|  |  | frequency |  |  |  |  |
|  |  | 8 kHz | 12 kHz | 16 kHz | 24 kHz | 32 kHz |
| P119S | +/+ (P36-38, n=6) vs. +/P119S (P36-39, n=6) | 0.9396 | 0.9567 | 0.8945 | 0.0866 | 0.4488 |
|  | +/+ (P36-38, n=6) vs. P119S/P119S (P19-20, n=6) | 0.8709 | <0.0001 | <0.0001 | <0.0001 | <0.0001 |
|  | +/+ (P36-38, n=6) vs. P119S/P119S (P36-39, n=6) | 0.0745 | <0.0001 | <0.0001 | <0.0001 | <0.0001 |
|  | +/P119S (P36-39, n=6) vs. P119S/P119S (P19-20, n=6) | 0.9975 | <0.0001 | <0.0001 | <0.0001 | <0.0001 |
|  | +/P119S (P36-39, n=6) vs. P119S/P119S (P36-39, n=6) | 0.2107 | <0.0001 | <0.0001 | <0.0001 | <0.0001 |
|  | P119S/P119S (P19-20, n=6) vs. P119S/P119S (P36-39, n=6) | 0.2851 | 0.0004 | 0.0003 | >0.9999 | 0.7176 |
|  |  | frequency |  |  |  |  |
|  |  | 8 kHz | 12 kHz | 16 kHz | 24 kHz | 32 kHz |
| A100T<br>P119S | +/+ (P36-40, n=10) vs. A100T/A100T (P20-21, n=8) | 0.3822 | <0.0001 | <0.0001 | <0.0001 | <0.0001 |
|  | +/+ (P36-40, n=10) vs. A100T/A100T (P30, n=6) | <0.0001 | <0.0001 | <0.0001 | <0.0001 | <0.0001 |
|  | A100T/A100T (P20-21, n=8) vs. A100T/A100T (P30, n=6) | <0.0001 | <0.0001 | <0.0001 | 0.9654 | 0.0273 |

